# Aging exerts a limited influence on the perception of self-generated and externally generated touch

**DOI:** 10.1101/2023.04.05.535662

**Authors:** Lili Timar, Xavier Job, Jean-Jacques Orban de Xivry, Konstantina Kilteni

## Abstract

Touch generated by our voluntary movements is attenuated both at the perceptual and neural level compared to touch of the same intensity delivered to our body by another person or machine. This somatosensory attenuation phenomenon relies on the integration of somatosensory input and predictions about the somatosensory consequences of our actions. Previous studies have reported increased somatosensory attenuation in elderly people, proposing an overreliance on sensorimotor predictions to compensate for age-related declines in somatosensory perception; however, recent results have challenged this direct relationship. In a preregistered study, we used a force-discrimination task to assess whether aging increases somatosensory attenuation and whether this increase is explained by decreased somatosensory precision in elderly individuals. Although 94% of our sample (n = 108, 21–77 years old) perceived their self-generated touches as weaker than externally generated touches of identical intensity (somatosensory attenuation) regardless of age, we did not find a significant increase in somatosensory attenuation in our elderly participants (65–77 years old), but a trend when considering only the oldest subset (69-77 years old). Moreover, we did not observe a significant age-related decline in somatosensory precision or a significant relationship of age with somatosensory attenuation. Together, our results suggest that aging exerts a limited influence on the perception of self-generated and externally generated touch and indicate a less direct relationship between somatosensory precision and attenuation in the elderly individuals than previously proposed.

**New and Noteworthy:** Self-generated touch is attenuated compared to externally generated touch of identical intensity. This somatosensory attenuation has been previously shown to be increased in elderly participants, but it remains unclear whether it is related to age-related somatosensory decline. In our preregistered study, we observed a trend for increased somatosensory attenuation in our oldest participants (≥69 years), but we found no evidence of an age-related decline in somatosensory function or a relationship of age with somatosensory attenuation.

## Introduction

Aging is associated with widespread brain changes (Raz et al. 2005; Zhao et al. 2019; Ziegler et al. 2012) that affect both motor (Michely et al. 2018; Solesio-Jofre et al. 2014; Wang et al. 2019; Zapparoli et al. 2022) and somatosensory systems (Brodoehl et al. 2013; Hagiwara et al. 2014; McIntyre et al. 2021; Wickremaratchi and Llewelyn 2006). In terms of motor performance, previous research has found that aging impairs the execution of voluntary movements (such as grasping), manual dexterity (such as grip force magnitude) (Diermayr et al. 2011), balance (Sturnieks et al. 2008), and motor learning (Vandevoorde and Orban de Xivry 2019; Vandevoorde and Orban De Xivry 2020; Wolpe et al. 2020). In addition, aging was shown to negatively influence somatosensory functioning, with multiple studies reporting an age-related decline (Bowden and McNulty 2013; Deflorio et al. 2022; Gescheider et al. 1994).

Motor control is largely dependent on the integration of motor signals with somatosensory information. A classic phenomenon related to this sensorimotor integration is somatosensory attenuation, which refers to perceiving touches that are produced by our own (voluntary) movements as less intense than touches of the same physical intensity that are externally generated (Bays and Wolpert 2008; Blakemore et al. 2000b; Kilteni 2023). For example, behavioral studies have shown that self-generated strokes, forces and taps applied to our left hand by our right hand are perceived as weaker than the same touches applied to our left hand by another person or a machine (Asimakidou et al. 2022; Bays et al. 2005, 2006; Blakemore et al. 1999a; Job and Kilteni 2023; Kilteni et al. 2018, 2019, 2020, 2021; Kilteni and Ehrsson 2017a, 2017b, 2020, 2022; Shergill et al. 2003). Similarly, neuroimaging studies have shown that self-generated touches elicit reduced activity in the primary (Hesse et al. 2010; Kilteni et al. 2023) and secondary somatosensory cortices (Blakemore et al. 1998; Kilteni and Ehrsson 2020; Shergill et al. 2013) as well as in the cerebellum (Blakemore et al. 1999b; Kilteni and Ehrsson 2020) compared to externally generated touches of identical intensity. Somatosensory attenuation is considered to facilitate differentiation between self-generated and externally generated sensations (Frith 2012) and to contribute to establishing and maintaining our sense of self by allowing us to separate our actions from those of others (Corlett et al. 2019; Frith 2005a). Furthermore, it is considered one of the reasons that humans are unable to tickle themselves (Blakemore et al. 2000b; Weiskrantz et al. 1971).

Computational motor control theories posit that somatosensory attenuation arises from the brain’s predictions about the sensory consequences of our movements. Accordingly, during a voluntary movement, the brain uses an internal forward model together with a copy of the motor command (“efference copy”) to predict the sensory feedback of the movement (Franklin and Wolpert 2011; Mcnamee and Wolpert 2019; Wolpert and Flanagan 2001). These predictions allow the brain to estimate the expected sensory feedback without relying on the actual sensory feedback, which suffers from intrinsic delays (Bays and Wolpert 2008; Davidson and Wolpert 2005; Franklin and Wolpert 2011; Kawato 1999; Shadmehr and Krakauer 2008), and to integrate it with the received sensory signals to improve the estimation of the state of the body (Shadmehr and Krakauer 2008). Action prediction signals also serve to attenuate the expected self-generated sensations (Bays et al. 2006; Job and Kilteni 2023), thereby increasing the salience and prioritizing the processing of unexpected externally generated sensations that might be more behaviorally relevant (Bays and Wolpert 2008; Blakemore et al. 2000b; Shergill et al. 2003; Wolpert and Flanagan 2001). Within a Bayesian integration framework, somatosensory attenuation relies on the integration of the forward model’s predictions and the somatosensory information, with both sources of information weighted based on their relative reliability (Ernst and Banks 2002; Körding et al. 2004). Interestingly, alterations in this integration of predictions and sensory information have been reported in several clinical and neurobiological models of psychosis spectrum disorders, such as schizophrenia (Blakemore et al. 2000a, 2002; Corlett et al. 2019; Frith 2005b, 2012; Frith et al. 2000; Shergill et al. 2005, 2014) and schizotypy (Asimakidou et al. 2022), as well as functional movement disorders (Pareés et al. 2014) and Parkinson’s disease (Wolpe et al. 2018).

Aberrant somatosensory attenuation has also been reported in elderly participants compared to young participants in two different studies (Parthasharathy et al. 2022; Wolpe et al. 2016). Specifically, when asked to match externally generated forces applied to their finger with self-produced forces, Wolpe et al. (2016) observed that older adults (65–88 years old) applied stronger self-produced forces than younger adults (18–39 years old), suggesting a greater attenuation of self-generated sensations with aging. Additionally, older adults were less precise than younger adults in distinguishing the different forces, indicating a negative impact of age on force sensitivity; the decreased force sensitivity was proportional to their increased attenuation. Based on these findings, the authors interpreted increased somatosensory attenuation in elderly individuals as decreased reliance on somatosensory information due to age-related reductions in somatosensory precision that, in turn, result in an increased reliance on sensorimotor predictions (consistent with Bayesian integration). On the other hand, Parthasharathy and colleagues (2022), using the same task but with the arm instead of the hand, also reported increased somatosensory attenuation in older adults (55–75 years old) compared to young adults (18–35 years old), similar to Wolpe et al. (2016), but found no evidence of decreased somatosensory precision in older adults, suggesting that somatosensory attenuation and precision might not be as closely related as previously suggested.

Here, we reinvestigated the role of aging in somatosensory attenuation and its relationship with somatosensory precision across a wide age range (21–77 years). Specifically, we tested whether a decline in somatosensory precision explains the effects of increased somatosensory attenuation with aging, as proposed by Wolpe et al. (2016), or if the two are unrelated, as suggested by Parthasarathy et al.(2022). The two previous studies used the force-matching task (Shergill et al. 2003) to quantify somatosensory attenuation, in which the participants receive an externally generated force on their relaxed left index finger by a motor and are subsequently asked to match this reference force. In the control condition, participants match the reference force by moving a joystick or slider that indirectly controls the force applied by the motor on their finger (slider condition). Several behavioral studies have shown that in this condition, participants precisely match the required forces, thus showing accurate somatosensory perception (Kilteni and Ehrsson 2017a, 2017b, 2020; Shergill et al. 2003; Wolpe et al. 2016). In contrast, in the experimental condition, when participants matched the reference force by directly pressing with their right index finger against their left one via a force sensor (direct condition), they overestimated the required forces and systematically produced stronger forces (Kilteni and Ehrsson 2020; Shergill et al. 2003; Wolpe et al. 2016). This suggests that participants attenuate their (directly) self-generated forces based on motor commands and increase the strength of self-produced forces to compensate for this somatosensory attenuation.

In the present study, we chose not to include the force-matching task and instead used the force-discrimination task, a well-established psychophysical test that has been previously used to assess somatosensory attenuation (Asimakidou et al. 2022; Bays et al. 2005, 2006; Job and Kilteni 2023; Kilteni 2023; Kilteni et al. 2019, 2020, 2021, 2023; Kilteni and Ehrsson 2022). In the force-discrimination task, participants receive two forces on their finger and are asked to indicate which force felt stronger. We chose the force-discrimination task instead of the force-matching task for three reasons. First, in contrast to the direct and slider conditions of the force-matching task, which require participants to move, the force-discrimination task allows a more accurate quantification of the perception of self-generated and externally generated forces because it includes a control condition of pure externally generated touch in the absence of any movement (no efference copy). Second, the force-discrimination task allows the psychophysical quantification of somatosensory precision for self-generated and externally generated stimuli separately. Third, elderly populations are known to have motor deficits, and their perception in the force-matching task is assessed with a motor response (*i.e.*, pressing to match a particular force or operating a joystick). Thus, another advantage of the force-discrimination task is that the perceptual report (*i.e.*, indicating which of two forces felt stronger) does not rely on motor abilities to the same extent. Moreover, given that both the force-matching task and the force-discrimination task involve the use of working memory to remember the forces to match (force-matching task) or judge them (force-discrimination task), we additionally assessed tactile working memory in our study for the first time to rule out the possibility that the increased somatosensory attenuation observed in older adults in the two previous studies was simply due to a decline in their tactile working memory.

## Materials and Methods

### Preregistration

The methods, hypotheses and analyses of the study were preregistered (https://osf.io/8u7by). All analyses included in the preregistration are indicated as “*preregistered analyses*” in the Results section. Any additional analyses that were not included in the preregistration are clearly indicated in the manuscript as “*supplementary analyses*” in the Results section.

### Participants

Data from one hundred and eight (108) participants were included in the present study. These participants were divided into the *young* (n = 36, age: range = 21–33; mean ± SD = 26 ± 3.85 years; 30 right-handed, 4 left-handed, 2 ambidextrous), *middle-aged* (n = 36, age: range = 43-56; mean ± SD = 48.6 ± 3.77 years; 30 right-handed, 3 left-handed, 3 ambidextrous) and *elderly* groups (n = 36, age: range = 65-77 years; mean ± SD = 69.6 ± 3.59 years; 35 right-handed, 1 left-handed). Each age group had a balanced sex ratio, consisting of 18 female and 18 male subjects. Handedness was assessed with the Edinburgh Handedness Inventory (Oldfield 1971). The sample size was based on a previous study assessing somatosensory attenuation and precision across similar age groups (Parthasharathy et al. 2022). All participants reported having normal or corrected-to-normal visual acuity, were healthy (without current or previous neurological or psychiatric disorders) and were not taking any medication to treat such conditions.

All participants provided written informed consent. The study lasted approximately 60 minutes and was approved by the Swedish Ethical Review Authority (application 2020-03186, amendment 2021-06235).

### Screening methods and exclusion criteria

#### Cognitive function

All elderly participants were tested for mild cognitive impairment, defined as greater cognitive impairment than is expected for one’s age. We used the Montreal Cognitive Assessment (MoCA version 8.3) (Nasreddine et al. 2005), which assesses cognitive function in several domains, including attention/working memory, executive function, episodic memory, language, and visuospatial skills; this assessment has been validated for use with individuals between 55 and 85 years old (Nasreddine et al. 2005). In the present study, the MoCA was used to screen elderly participants and ensure that they could understand and follow experimental instructions. Scoring of each individual and correction for low education level were performed according to the instructions. The test was conducted in the native language of the participant by a certified experimenter who completed the necessary training to carry out and score the test (https://www.mocatest.org/training-certification/). Following the official MoCA scoring instructions regarding the cut-off score, we included only elderly individuals with a MoCA score of 26 or higher.

#### Tactile working memory

All participants were assessed for tactile working memory (WM) to ensure that they could reliably remember at least two brief forces applied to their fingers in a short period of time, as required by the force-discrimination task (see below). We used the working memory task introduced and described by Heled et al. (2021). During the task, the participants comfortably sat in a chair with their eyes closed and placed four fingers of each hand on the upper row of a QWERTY keyboard (right hand fingers on ‘Q’, ‘W’, ‘E’, ‘R’ keys and left hand fingers on ‘U’, ‘I’, ‘O’, ‘P’ keys). Next, the experimenter lightly touched the participant’s fingers, between the second and third knuckle, with the back of a pencil for one second each, in a specific sequence. Participants were then asked to repeat the sequence back, in the same order as it was presented, by pushing down on the keys with the fingers that had been touched (**Supplementary Figure S1**). One elderly participant had difficulties with the keyboard, and he provided the answers verbally by naming the fingers instead of tapping on the keys. The test started with three 2-finger sequence trials. If at least one of the three sequences was correctly reproduced, then the next sequence was increased in length by one, up to sequences 9 fingers in length. Each sequence length included three trials: one trial on the left hand only, one on the right hand only, and one on both hands. The task ended if participant made three consecutive mistakes within the same sequence length or when the ninth sequence was successfully recalled. We calculated the longest sequence that the participant could recall without a mistake (*longest sequence recalled*; score range: 0–9) and the number of correct answers given (*maximum WM score*; score range: 0–24). We included individuals who could recall sequences of at least two fingers (*longest sequence recalled* ≥ 2), given that the force-discrimination task included two tactile stimuli.

#### Exclusion of participants

In total, eighteen (18) participants were excluded: fourteen elderly participants who did not reach the MoCA cutoff score, one middle-aged participant who could not perform the working memory task, one middle-aged and one elderly participant who experienced technical issues, and finally, one middle-aged participant who revealed that they took medication after being tested. These excluded individuals were replaced by an equal number of new participants to reach the target sample size (108).

### Psychophysical task

The psychophysical task was a two-alternative forced choice (2AFC) force-discrimination task that has been used by numerous studies investigating somatosensory attenuation (Asimakidou et al. 2022; Bays et al. 2005, 2006; Job and Kilteni 2023; Kilteni 2023; Kilteni et al. 2020, 2021, 2023; Kilteni and Ehrsson 2022).

Participants rested their left hands palm up with their index fingers on a molded support and their right hands palm down on top of a set of sponges. A vacuum pillow (Germa protec, AB Germa) was provided to support the participants’ left arm and increase their comfort. Every trial started with an auditory tone. Next, a DC electric motor (Maxon EC Motor EC 90 flat; manufactured in Switzerland) delivered two brief (100-ms) forces to the pulp of participants’ left index finger through a cylindrical probe (25 mm in height) with a flat aluminum surface (20 mm in diameter) attached to the motor’s lever. Participants then verbally indicated which force felt stronger, the first (*test* force) or the second (*comparison* force). The interstimulus interval varied randomly between 500 ms and 800 ms. The intensity of the *test* force was set to 2 N, while the *comparison* force pseudorandomly varied among seven possible intensities (1, 1.5, 1.75, 2, 2.25, 2.5, or 3 N). A force sensor (FSG15N1A, Honeywell Inc.; diameter, 5 mm; minimum resolution, 0.01 N; response time, 1 ms; measurement range, 0–15 N) was placed within the cylindrical probe to record the forces exerted on the left index finger. A force of 0.1 N was constantly applied to the participant’s left index finger to ensure accurate force intensities.

There were two experimental conditions. In the *externally generated touch* condition (**Figure 1a**), the participants relaxed both their hands, and the *test* force was delivered automatically 800 ms after the auditory tone. In the *self-generated touch* condition (**Figure 1b**), the participants were instructed to tap with their right index finger on a force sensor (identical specifications as above) placed on top of, but not in contact with, their left index finger. The participants’ tap on the force sensor triggered the *test* force on their left index finger. Each condition consisted of 70 trials; all seven intensities of the *comparison* force were presented ten times (7×10) per condition, resulting in a total of 140 trials per participant. The order of the intensities was pseudorandomized, and the order of the conditions was counterbalanced across participants.

**Figure 1.**
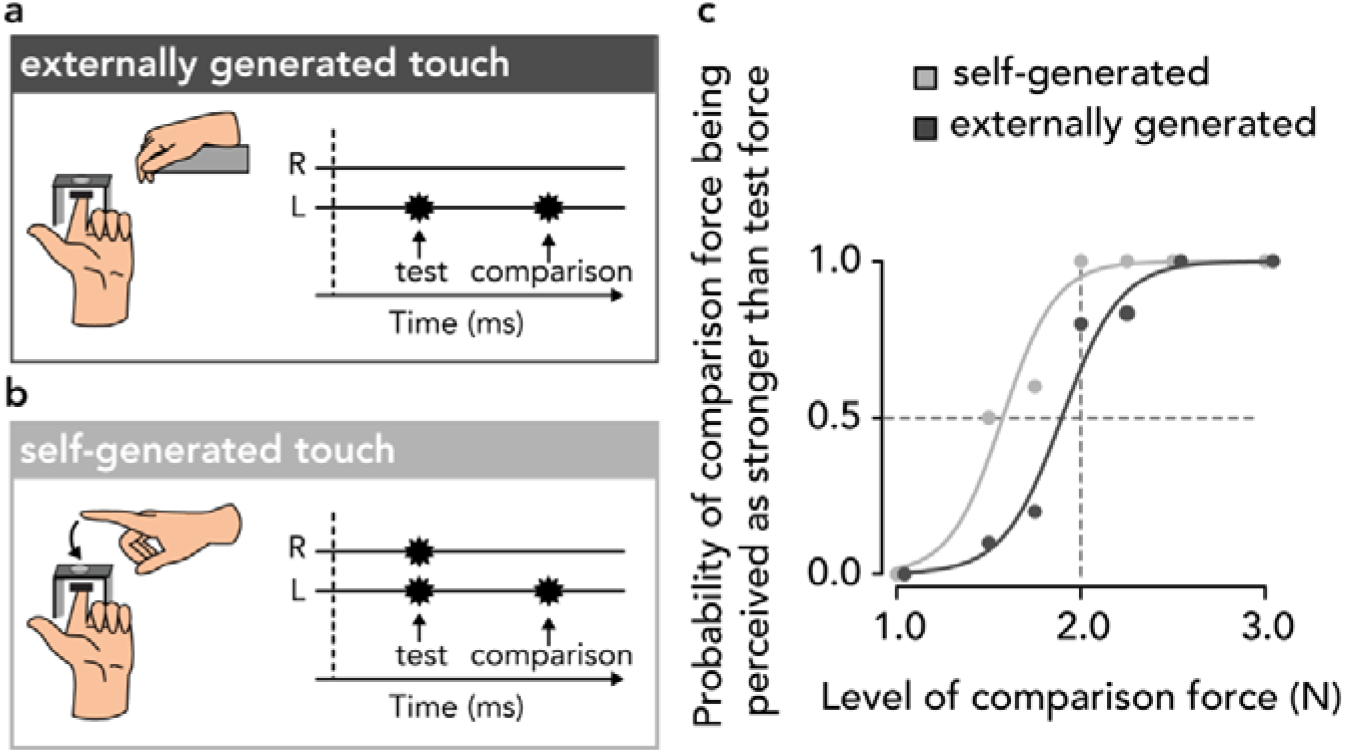
The force-discrimination task. In both conditions, the participants experienced two forces on the pulp of their left index finger, the *test* force and the *comparison* force, and verbally indicated which force felt stronger. **(a)** In the *externally generated touch* condition, the participants relaxed both their hands and received the *test* and the *comparison* forces automatically on the pulp of their left index finger. **(b)** In the *self-generated touch* condition, the participants triggered the *test* force on the left index finger by actively tapping on a force sensor with their right index finger placed above their left finger. Next, they received the *comparison* force. **(c)** Responses and fitted logistic models of the responses of one participant in the two experimental conditions. The leftward shift of the light gray curve with respect to the dark gray one indicates that the *test* force in the *self-generated touch* condition felt weaker compared to that in the *externally generated touch* condition.

White noise was played through a pair of headphones to mask any sounds made by the motor. During the experiment, the participants’ left index finger was occluded from vision, and they were asked to focus on a fixation cross placed on the wall approximately 80 cm in front of them.

### Psychophysical fit

In each condition, the participant’s responses were fitted with a generalized linear model using a *logit* link function implemented in R (version 4.2.0) (R Core Team 2022) within the function *glm* with the default fitting method (reweighted least squares) (**Figure 1c**) (Equation 1):

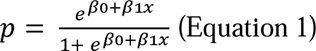

Two parameters of interest were extracted. First, the point of subjective equality 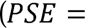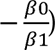 represents the intensity at which the *test* force felt as strong as the *comparison* force (*p* = 0.5) and thus quantifies the participants’ perceived intensity of the *test* force. Subsequently, somatosensory attenuation was calculated as the difference between the PSEs of the *externally generated* and *self-generated* touch conditions (*PSE_external_* – *PSE_self_*) (Asimakidou et al. 2022; Job and Kilteni 2023; Kilteni et al. 2020, 2023). That is, a participant shows somatosensory attenuation if their *PSE_self_* is smaller than the *PSE_external_*. Second, the *just noticeable difference* 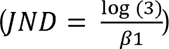 reflects the participants’ sensitivity in the psychophysical task and thus quantifies their somatosensory precision in each condition, corresponding to the difference between the thresholds at *p* = 0.5 and *p* = 0.75.

Before fitting the responses, the *comparison* forces were binned to the nearest of the seven possible force intensities (1, 1.5, 1.75, 2, 2.25, 2.5, or 3 N). After the data collection, 60 out of 15120 (0.4%) trials were rejected: 42 trials (0.28%) were rejected because the intensity of the *test* force (2 N) was not applied accurately (*test* force < 1.85 N or *test* force > 2.15 N), and 18 trials (0.12%) were rejected because there were missing responses.

### Additional measures

As secondary variables, we further recorded (a) the peak active forces the participants applied to the force sensor with their right index finger (*peak force*), (b) the time it took for the participants to reach the peak force after the beginning of the trial (*time to peak force*), and (c) the movements of their right index finger as registered using a Micro Sensor 1.8 attached to a Polhemus Liberty electromagnetic tracker (https://polhemus.com/motion-tracking/all-trackers/liberty). If somatosensory attenuation is increased in *elderly* participants compared to *younger* participants, as we expected, these additional measures could be used to explore the relationships of age with forces, timing, and kinematics together with attenuation. Due to technical reasons, the movements of the right index finger were not correctly registered; thus, supplementary analyses were performed with only the active peak forces and their times.

### Hypotheses

We tested four preregistered experimental hypotheses using the collected data. First, we expected to replicate the classic somatosensory attenuation phenomenon in our sample by finding that the PSEs in the *self-generated touch* condition were significantly lower than the PSEs in the *externally generated touch* condition, regardless of age group (H1). Second, given earlier studies reporting a decline in somatosensory functioning (Bowden and McNulty 2013; Deflorio et al. 2022; Gescheider et al. 1994; Humes et al. 2009; Stevens and Cruz 1996) and a reduction in the density of cutaneous mechanoreceptors with age (García-Piqueras et al. 2019) (see also (Lin et al. 2004)), we hypothesized that JND values in the *externally generated touch* condition (*i.e.*, *JND_external_*) would be significantly higher in *elderly* participants than in *young* and *middle-aged* participants (H2). Third, given the two previous studies reporting increased attenuation in older participants (Parthasharathy et al. 2022; Wolpe et al. 2016), we expected to find increased somatosensory attenuation in *elderly* participants compared with *younger* participants (H3). Finally, we assessed the proposal of Wolpe et al. (2016) that decreased somatosensory precision drives the increased attenuation in *elderly* participants by testing whether somatosensory precision is a significant positive predictor of somatosensory attenuation (H4).

### Statistical analysis

Data were analyzed in R (version 4.2.0) (R Core Team 2022) and JASP (version 0.16.4) (JASP Team 2022). The normality of the data was assessed with the Shapiro[Wilk test. Planned comparisons were performed using parametric (paired or independent-sample *t* tests) or nonparametric (Wilcoxon signed-rank and Wilcoxon rank sum) tests depending on the normality of variable distributions. A Welch *t* test was used if the variances of the compared distributions were unequal according to Levene’s test. For every statistical comparison, we report the corresponding statistic, the 95% confidence intervals (*CI^95^*) and the effect size (Cohen’s *d* or the matched rank-biserial correlation (*r_rb_*), depending on the distribution normality). We also performed a Bayesian factor (*BF_01_*) analysis (default Cauchy priors with a scale of 0.707) for the statistical tests of interest reporting nonsignificant differences to provide information about the level of support for the null hypothesis compared to the alternative hypothesis. For correlation analyses, we computed multilevel correlations to account for differences between the age groups. We report Spearman’s *rho* correlation coefficients for non-normally distributed data. Bayes factors (*BF_01_*) are provided for the nonsignificant correlation tests too. Finally, for regression analysis, a robust linear regression was performed to reduce the impact of outlier observations.

Two-tailed statistical tests were used to test all four preregistered (https://osf.io/8u7by) and supplementary hypotheses. When performing multiple comparisons among the three age groups, we corrected the *p* values using the false discovery rate (*FDR*). Corrected p values are thus denoted as “*FDR-corrected*” throughout.

## Results

As stated in our inclusion criteria, we first ensured that our elderly participants showed no signs of mild cognitive impairment and that all participants could retain at least two tactile stimuli applied to their fingers in their working memory (**Supplementary Text S1**, **Figure S2**).

### Somatosensory attenuation – preregistered analysis

Our first hypothesis was that PSEs in the *self-generated touch* condition would be significantly lower than the PSEs in the *externally generated touch* condition, regardless of age group. Supporting our first hypothesis (H1), the PSEs in the *self-generated touch* condition were significantly lower than those in the *externally generated touch* condition across the entire sample (*n* = 108): *W* = 112, *p* < .001, *CI^95^* = [-0.310, −0.224], *rrb* = −0.962 (**Figure 2a, Supplementary Figures S3-S5**). This pattern was observed in 102 out of 108 (94%) participants and indicates that self-generated forces are robustly attenuated compared to externally generated forces of equal intensity, in line with several previous studies (Asimakidou et al. 2022; Bays et al. 2005, 2006; Job and Kilteni 2023; Kilteni et al. 2019, 2020, 2021, 2023).

**Figure 2.**
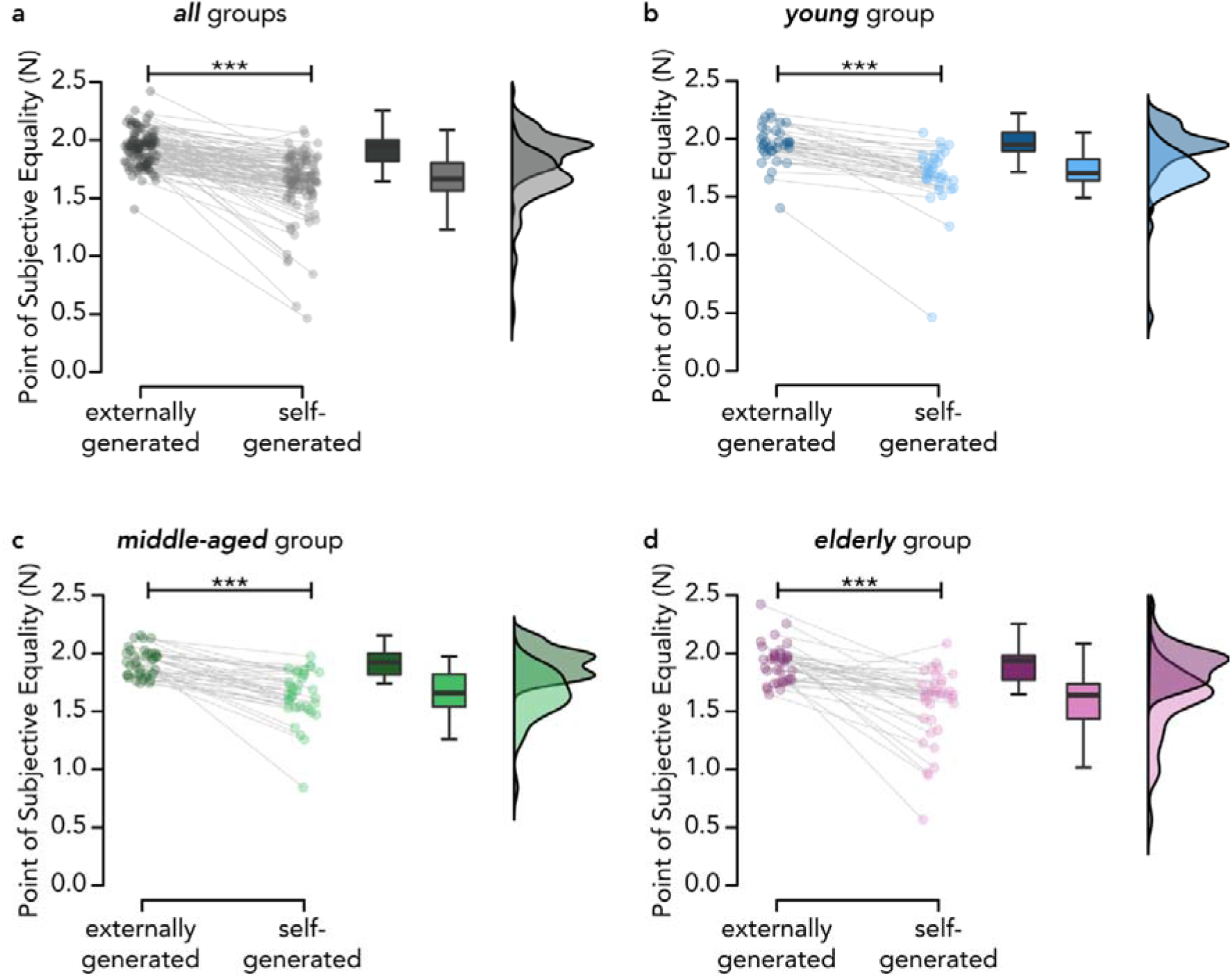
Somatosensory attenuation across age groups. **(a)** Across all age groups (pooled data; n = 108), self-generated touches were perceived as significantly weaker than externally generated touches of identical intensity. The same effect was found separately for the *young* **(b)**, *middle-aged* **(c)**, and *elderly* groups **(d)** (n = 36 for each group). The boxplots display the median and interquartile ranges of the PSEs in the *externally generated* and *self-generated touch* conditions per age group. Markers denote the PSE values for each participant, and raincloud plots show the distribution of the data. Line plots illustrate the PSE differences between the *externally generated* and *self-generated touch* conditions for each participant (*** *p* <.001).

### Somatosensory attenuation – supplementary analysis

Additional supplementary analyses showed that the attenuation effect was observed in every age group: PSEs in the *self-generated touch* condition were significantly lower than those in the *externally generated touch* condition within the *young* (*W* = 0, *p* < .001, *CI^95^* = [-0.289, - 0.193], *rrb* = −1.0) (**Figure 2b**), *middle-aged* (*W* = 10, *p* < .001, *CI^95^* = [-0.329, −0.194], *rrb* = −0.970) (**Figure 2c**) and *elderly* group (*W* = 26, *p* < .001, *CI^95^*= [-0.446, −0.210], *rrb* = - 0.922) (**Figure 2d**).

### Aging and somatosensory precision – preregistered analysis

Second, we hypothesized that JND values in the *externally generated touch* condition (*i.e.*, *JND_external_*) would be significantly higher for *elderly* participants than for *young* and *middle-aged* participants. Contrary to our hypothesis (H2), we did not find an increase in the JND values in the *elderly* group compared to the *young* group (*W* = 514, *p* = 0.399 *FDR*-corrected, *CI^95^* = [-0.053, 0.008], *rrb* = −0.207). The Bayesian analysis provided anecdotal support for the absence of a difference in somatosensory precision between the *elderly* and *young* groups (*BF_01_*= 1.549). No differences were observed between the *elderly* and *middle-aged* groups (*W* = 564, *p* = 0.468 *FDR*-corrected, *CI^95^*= [-0.040, 0.017], *rrb* = −0.130) and the Bayesian analysis provided moderate support for the absence of difference (*BF_01_* = 3.210). Finally, the JND values of the *middle-aged* group did not significantly differ from those of the *young* group (*W* = 583, *p* = 0.468 *FDR*-corrected, *CI^95^* = [-0.038, 0.016], *rrb* = −0.100, *BF_01_* = 2.824) (**Figure 3**).

**Figure 3.**
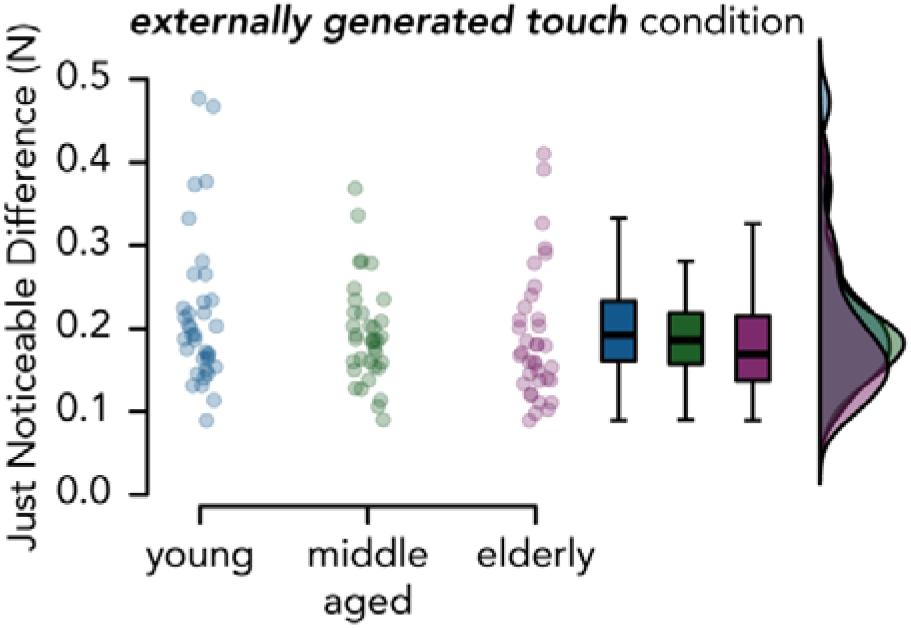
Somatosensory precision across age groups. JND values in the *externally generated touch* condition across the three age groups. There were no significant differences among the three groups, and the Bayesian analyses supported the absence of differences. The boxplots display the median and interquartile ranges, and the dots represent the individual participant values. Raincloud plots show the distribution of the data.

### Aging and somatosensory precision – supplementary analysis

In a non-preregistered (supplementary) post hoc analysis, we explored whether somatosensory impairment was more pronounced in the oldest of our *elderly* participants. To this end, we performed the same analysis as above, but we split the *elderly* group (65–77 years of age) at its median age and compared the oldest *elderly 69+* participants (*n* = 18, *age* = 69–77 years) to the *young* group. Once again, we did not detect any somatosensory impairment in the *elderly 69+* participants compared to the *young* participants (*W* = 220, *p* = 0.058, *CI^95^* = [-0.064, 0.003], *rrb* = −0.321, *BF_01_* = 1.207) (**Supplementary Figure S6**). If anything, the pattern suggested similar if not better somatosensory precision in the *elderly 69+* participants compared to the *young* participants.

### Aging and somatosensory attenuation – preregistered analysis

To test our third hypothesis, we examined whether the magnitude of somatosensory attenuation was greater in the *elderly* group than in the other two groups, as previously shown (Parthasharathy et al. 2022; Wolpe et al. 2016). Contrary to our hypothesis (H3), we did not observe any significant increase in the magnitude of somatosensory attenuation between the *elderly* group and the *young* (*W* = 710, *p* = 0.736 *FDR*-corrected, *CI^95^* = [-0.049, 0.149], *rrb* = 0.096) or the *middle-aged* group (*W* = 712, *p* = 0.736 *FDR*-corrected, *CI^95^* = [-0.077, 0.159], *rrb* = 0.099), nor between the *middle-aged* and *young* groups (*W* = 673, *p* = 0.784 *FDR*-corrected, *CI^95^*= [-0.069, 0.095], *rrb* = 0.039) (**Figure 4a**). The Bayesian analysis provided moderate evidence of a similar magnitude of attenuation across all three age groups (*elderly* compared to *young*: *BF_01_*= 3.732; *elderly* compared to *middle-aged*: *BF_01_*= 3.206; *middle-aged* compared to *young*: *BF_01_*= 4.139).

**Figure 4.**
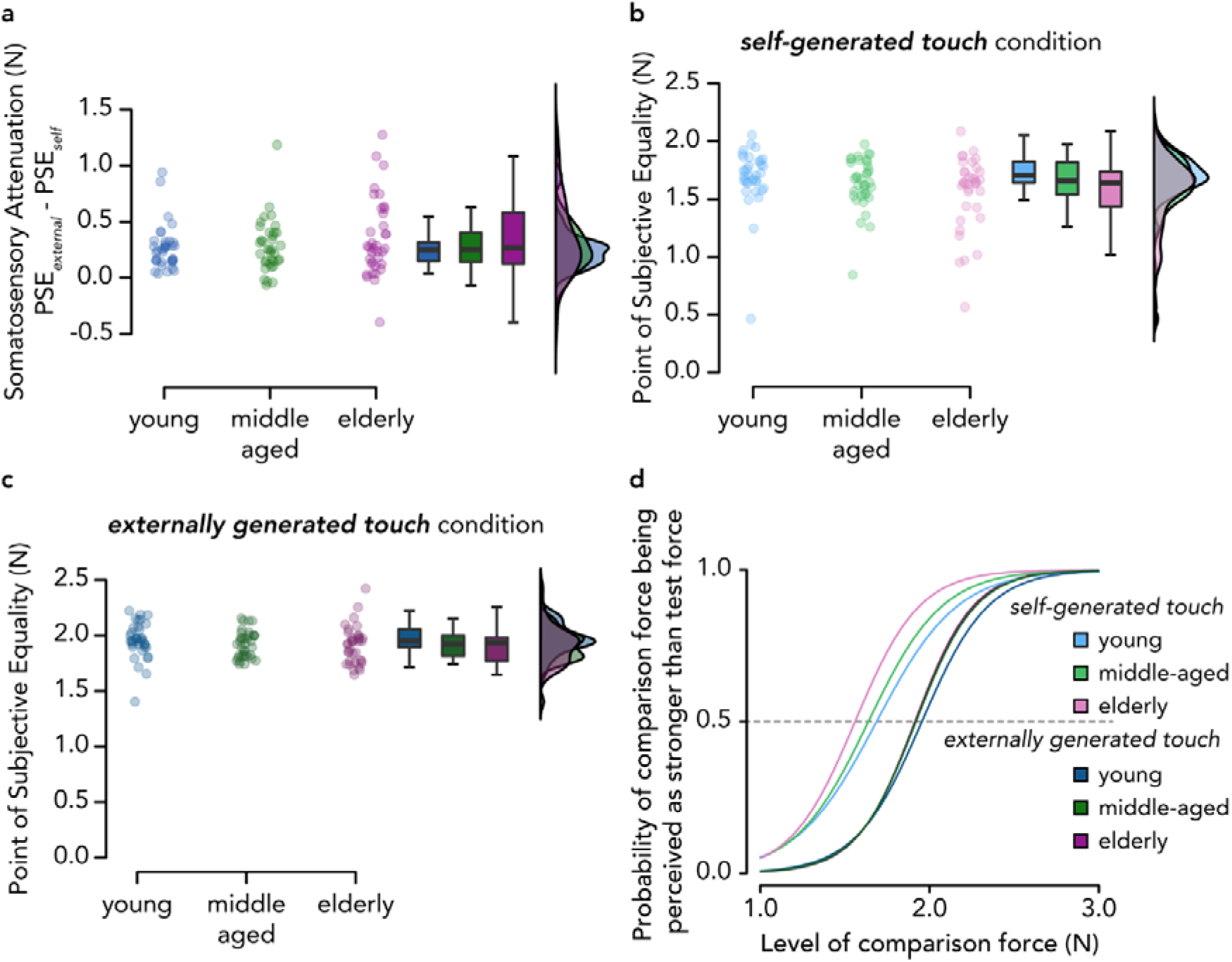
Somatosensory attenuation across age groups. **(a)** Somatosensory attenuation (*PSE_external_* – *PSE_self_*) across the three age groups. No significant increase in somatosensory attenuation was observed in the *elderly* group compared to the *middle-aged* and *young* groups or between the *middle-aged* group and the *young* group. (**b-c**) The *elderly* group showed a tendency to perceive their self-generated touches as weaker compared to the *young* group (**b**), and a similar tendency was observed for externally generated touches (**c**), indicating weaker somatosensory perception in elderly participants in general. (**d**) Mean psychometric curves for each age group and experimental condition according to the mean PSE and JND values. A leftward shift of the curve in the *self-generated touch* condition compared to the *externally generated touch* condition indicates somatosensory attenuation. The curves for the *externally generated touch* condition overlap for the *middle-aged* and *elderly* participants.

### Aging and somatosensory attenuation – supplementary analyses

First, to further explore this absence of increased attenuation in the *elderly* participants, we performed two additional non-preregistered analyses to test whether the participants’ perception differed in the *self-generated* and *externally generated touch* conditions within each age group. As seen in the boxplots of **Figure 4b-c** and the group model fits in **Figure 4d**, the PSEs in both the *self-generated touch* and *externally generated touch* conditions decreased as a function of aging, which could effectively explain why we did not observe significant changes in the magnitude of somatosensory attenuation (*i.e.*, no PSE difference between the two conditions).

However, there were no significant differences among groups in either the *self-generated touch* condition (*elderly* vs. *young* group, *W* = 456.5, *p* = 0.093 *FDR-corrected*, *CI^95^* = [- 0.195, −0.007], *rrb* = −0.296, *BF_01_* = 0.734; *elderly* vs. *middle-aged*, *W* = 586, *p* = 0.489 *FDR-corrected*, *CI^95^* = [-0.155, 0.057], *rrb* = −0.096, *BF_01_* = 3.170; *middle-aged* vs. *young, W* = 527.5, *p* = 0.265 *FDR-corrected*, *CI^95^* = [-0.137, 0.025], *rrb* = −0.186, *BF_01_* = 1.825), or the *externally generated touch* condition (*elderly* vs. *young* group, *W* = 519.5, *p* = 0.308 *FDR-corrected*, *CI^95^* = [-0.133, 0.024], *rrb* = −0.198, *BF_01_* = 2.011; *elderly* vs. *middle-aged, t*(70) = −0.153, *p* = 0.879 *FDR-corrected*, *CI^95^* = [-0.073, 0.063], *d* = −0.036, *BF_01_* = 4.073; *middle-aged* vs. *young*, *W* = 535, *p* = 0.308 *FDR-corrected*, *CI^95^* = [-0.122, 0.025], *rrb* = −0.174, *BF_01_* = 2.015).

Second, we performed the same non-preregistered post hoc analysis used to test Hypothesis 2 for Hypothesis 3 to assess whether increased somatosensory attenuation would be more pronounced in the oldest of our *elderly* participants. As before, we split the elderly group (age range: 65–77 years) at the median age of our elderly participants, and we compared the *elderly 69+* participants (*n* = 18, *age* = 69–77 years) to the *young* group. Indeed, we observed a trend for somatosensory attenuation being higher in the *elderly 69+* compared to the *young* group (*W* = 415, *p* = 0.097, *CI^95^* = [-0.017, 0.334], *rrb* = 0.281, *BF_01_* = 1.885) (**Figure 5**).

**Figure 5.**
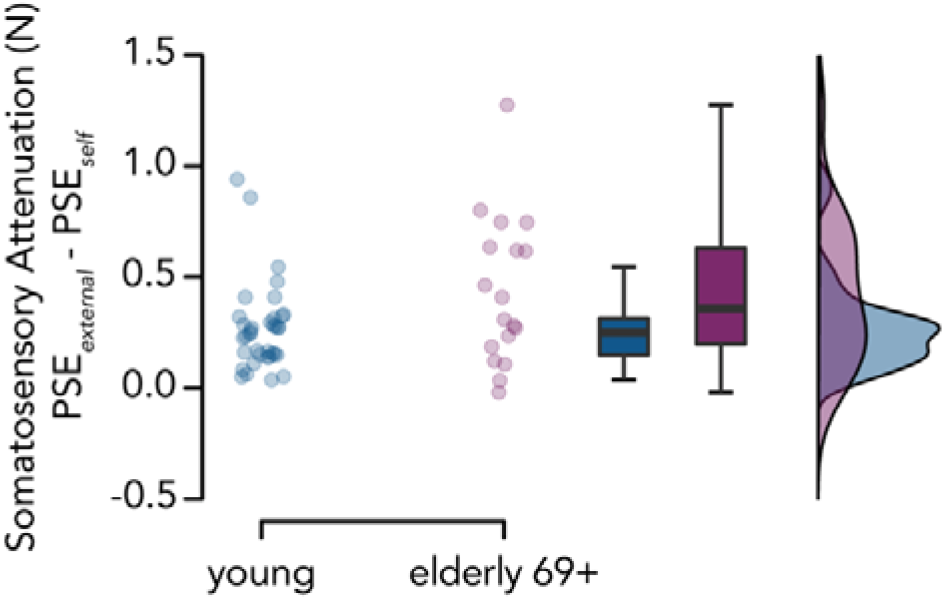
Somatosensory attenuation in *young* and *elderly 69+* participants. We observed a tendency for greater somatosensory attenuation in the *elderly* 69+ group (*n* = 18) than in the *young* group (*n* = 36) which did not reach statistical significance.

### Somatosensory attenuation, aging, and somatosensory precision – preregistered analysis

Finally, to test our fourth and final hypothesis, we investigated whether the magnitude of somatosensory attenuation is related to the somatosensory precision of externally generated touch by testing whether somatosensory precision is a significant positive predictor of somatosensory attenuation, as previously suggested (Wolpe et al. 2016). To this end, we constructed a robust linear regression model using somatosensory precision as a regressor of somatosensory attenuation as well as age group (*young*, *middle-aged*, *elderly*) and their interaction. We chose a robust linear regression model rather than a linear regression model to decrease the effect of outliers. None of the regressor coefficients or their interaction were significant (all *p* values > 0.700, *R^2^*= 0.010). In line with our above results, somatosensory precision was not a predictor of somatosensory attenuation, and somatosensory precision and age did not exert a joint effect on the degree of somatosensory attenuation.

### Additional measures

Finally, there were no significant differences in the magnitude of the active forces the participants applied or in the time it took them to apply the forces among age groups, and there was no significant relationship between these measures and somatosensory attenuation (**Supplementary Text S2, Supplementary Figure S7)**. Similarly, no significant relationship was detected between tactile working memory and somatosensory attenuation **(Supplementary Text S3**).

## Discussion

The present study investigated how aging impacts somatosensory attenuation and somatosensory precision, with the aim of resolving previous contradictory results regarding the underlying mechanisms of age-related changes in somatosensory attenuation (Parthasharathy et al. 2022; Wolpe et al. 2016).

Our first analysis replicated the somatosensory attenuation phenomenon across our entire sample. Specifically, the prevalence of somatosensory attenuation was high (94% of the 108 participants showed this effect), in line with studies using similar (Asimakidou et al. 2022) or larger sample sizes (Wolpe et al. 2016). The attenuation effect was detected in each individual age group: *young*, *middle-aged,* and *elderly* participants exhibited significant somatosensory attenuation of self-generated forces compared to externally generated forces of the same intensity. These results therefore extend those of earlier studies (Parthasharathy et al. 2022; Wolpe et al. 2016), including the use of the force-discrimination task to psychophysically quantify somatosensory attenuation across different age groups.

Contrary to our hypothesis and to previous evidence showing a decline in somatosensory precision with aging (Bowden and McNulty 2013; Deflorio et al. 2022; Gescheider et al. 1994; Humes et al. 2009; Stevens and Cruz 1996), we did not find that *elderly* participants were worse than *young* participants in discriminating forces. Furthermore, this absence of decline was supported by our Bayesian analysis. Although this result is surprising, several factors could account for this lack of somatosensory decline with aging. First, it could be argued that our psychophysical task (force-discrimination) was not sensitive enough to capture potentially small differences in precision among age groups. However, we consider this unlikely since we have previously used this task to detect differences in somatosensory precision (Kilteni and Ehrsson 2022); moreover, as shown in **Figure 3** and **Figure S6**, our *elderly* participants demonstrated (albeit not significantly) better performance than the *young* participants, as also found by Parthasharathy et al. (2022). Second, our participants may not have been old enough to manifest somatosensory deficits. For example, Bowden and McNulty (2013) showed significantly elevated tactile thresholds at the tip of the index finger for only adults above 80 years old. Moreover, by combining different tests of somatosensory function, these authors concluded that the decline in cutaneous sensation becomes faster after the age of 60 years in males and 70 years in females. However, we also consider this interpretation unlikely, as we did not observe somatosensory deficits even when comparing the *young* group to the oldest subset of participants from our *elderly* group (individuals ≥ 69 years old), who were predominantly male and thus should have exhibited greater somatosensory deficits. However, in addition to their chronological age, we should mention that all our *elderly* participants were screened to prevent the presence of mild cognitive decline. Since sensorimotor and cognitive deficits are comorbid in older adults, and cognitive decline is linked with deficits in sensory function (Ghisletta and Lindenberger 2005; Li and Lindenberger 2002; Lindenberger and Baltes 1994; Roberts and Allen 2016; Rong et al. 2020), one possibility is that our screened elderly sample was skewed toward individuals with better cognitive and sensory abilities than the elderly samples of previous studies. Relatedly, another possibility is that our older sample might have used remaining intact cognitive processes to compensate for any age-related somatosensory decline and perform at a similar level as younger adults (Roberts and Allen 2016).

An alternative explanation for the lack of somatosensory deficits with aging could be that somatosensory decline is minimal and/or not always present in elderly participants (Heft and Robinson 2017). It is interesting to note that age-related somatosensory deficits are less systematically reported than visual or auditory deficits (Heft and Robinson 2014, 2017), do not necessarily co-occur with deficits in other sensory modalities (Cavazzana et al. 2018), and can highly depend on the sex of the participants, the stimulation site and assessment method (Bowden and McNulty 2013). In contrast to studies reporting somatosensory decline, other studies report minimal or even no somatosensory changes between young and elderly participants. For example, in a fine texture-discrimination task, Skedung et al. (2018) reported lower discrimination capacity in the elderly group (aged 67–85 years) than the young group (aged 19–25 years), with 13 out of 30 elderly participants (43%) nevertheless performing equally as well as the young participants. Older participants (mean age = 63 years) were shown to have similar haptic thresholds for detection and discrimination as younger participants (mean age = 28 years) (Konczak et al. 2012), and chronological age (50–100 years) was not found to significantly correlate with tactile measures (Cavazzana et al. 2018). Additionally, in a pressure sensitivity task, Tremblay et al. (2005) observed that older (60–86 years) participants’ sensitivity to minimal pressure was highly functional, even if it was reduced compared to that of younger participants (aged 19–32 years). Similar to our results, Parthasharathy et al. (2022) reported that older participants reproduced the forces more accurately in the slider condition of the force-matching task than young participants. Overall, it could be that somatosensory function shows minimal to small declines with age (Heft and Robinson 2014), similar to proprioception, which shows a small, if nonnegligible, age-related decline (Djajadikarta et al. 2020; Herter et al. 2014; Kitchen and Miall 2021; Roberts and Allen 2016).

Finally, it is also possible that pressure/force perception in elderly individuals is more resistant to age-related decline than other types of tactile functioning. Interestingly, most of the studies showing large declines in somatosensory sensitivity with aging used texture discrimination, spatial acuity or vibrotactile tasks (Gescheider et al. 1994; Skedung et al. 2018; Stevens and Cruz 1996), but less consistent findings were shown for pressure/force perception (Parthasharathy et al. 2022; Tremblay et al. 2005; Wolpe et al. 2016). This might not be surprising, as different assessments of somatosensory functioning might stimulate distinct classes of mechanoreceptors that may be differentially affected by aging (García-Piqueras et al. 2019).

In contrast to our hypothesis, we did not find significantly higher somatosensory attenuation in the elderly group than in the younger groups, as reported by Wolpe et al. (2016) and Parthasharathy (2022). Although, as seen in **Figure 4b**, *elderly* participants tended to perceive their self-generated touches as weaker than younger participants, the same pattern was observed for externally generated touches (**Figure 4c**). We speculate that increased attenuation might be pronounced in the oldest of our participants, as Wolpe et al. (2016) found a sharp increase in attenuation at the higher end of their age group, suggesting a rapid increase in the attenuation of self-generated forces in individuals in their late 70s and 80 years or older, rather than a linear relationship with age. Indeed, when we compared the oldest subset of our elderly participants (aged ≥69 years) to the *young* group, we did observe a tendency for higher attenuation that follows the same pattern as previous studies (Parthasharathy et al. 2022; Wolpe et al. 2016), albeit not statistically significant. This suggests that the increase in somatosensory attenuation might require older samples than previously suggested.

It is important to mention that the current study employed the two-alternative forced choice force-discrimination task to measure somatosensory attenuation, while the previous ones used the force-matching task (Parthasharathy et al. 2022; Wolpe et al. 2016). The force-discrimination task has the advantage not to rely on motor abilities when participants give their perceptual responses. This contrasts with the force-matching task where the participants need to press or move the slider to give their response. Given that elderly participants often show deteriorated motor performance compared to young ones (*e.g.*, (Frolov et al. 2020; Zapparoli et al. 2022)), using the force-discrimination task should theoretically remove such confounds when assessing somatosensory attenuation. Moreover, the force-discrimination task directly quantifies the somatosensory precision and perceived magnitude of externally generated forces given that it includes a condition of externally generated forces (*i.e.*, in the absence of a motor command/efference copy). This contrasts the force-matching task that assesses the participants’ baseline somatosensory perception via the slider condition that requires them to move (*i.e.*, motor command). To this end, it could be argued that the force-discrimination task assessed somatosensory attenuation in elderly participants more directly than the force-matching task.

However, the force-discrimination task also requires that participants keep two forces in their working memory and compare their intensity, while the force-matching task requires remembering the intensity of a single force. Additionally, the force-discrimination task we used in the present study included 70 trials per condition while the force-matching task, as in the study of Wolpe et al. (2016), included only 32 trials. Hence, it could be argued that the force-discrimination task requires larger working memory capacity and is more taxing on the participants’ cognitive abilities than the force-matching task – a difference that would be more prevalent when comparing elderly with young participants with known differences in their working memory and cognitive abilities. However, we consider it unlikely that our absence of increased somatosensory attenuation effects in elderly is driven by the potential increased working memory demands of the force-discrimination task. First, we carefully screened our participants, and we allowed only the ones that could show no mild cognitive impairment and keep at least two forces in their memory according to a validated standardized tactile working memory task (Heled et al., 2021) to participate. Second, we found no relationship between working memory and the magnitude of somatosensory attenuation in our supplementary analyses, and the Bayesian analysis supported the absence of such relationship (**Supplementary Text S3**). Third, despite being more in number, the trials in the force-discrimination task are shorter than the trials in the force-matching task because the forces participants receive, albeit two, are substantially shorter (*e.g.*, 100 ms duration each with 500-800 ms interval between them) than the typical duration of the reference force in the force-matching task (*e.g.*, 3 seconds). Executing a single condition of the force-discrimination task took approximately ∼10 minutes. Importantly, if working memory or cognitive abilities were driving the performance in the force-discrimination task, we would expect to find significant differences between the *young* and the *elderly* group in the experimental conditions - which we did not.

Finally, across our sample, we did not find any significant relationship between somatosensory attenuation and somatosensory precision. This is in agreement with our previous findings reporting no significant relationship between perceived somatosensory precision and somatosensory magnitude (Kilteni and Ehrsson 2022). According to the Bayesian integration framework, the age-related increase in somatosensory attenuation is caused by increased weighting of the internal models’ predictions and decreased weighting of sensory information (Wolpe et al. 2016). Given that the internal model is thought to remain intact with aging (Heuer et al. 2011; Vandevoorde and Orban de Xivry 2019), somatosensory decline should lead to increased somatosensory attenuation. Since we did not observe any somatosensory decline in our *elderly* participants, we might not have recruited a sample with enough variability to detect such a relationship. Nevertheless, since we observed a tendency for higher somatosensory attenuation in our oldest participants without a similar tendency for concomitant somatosensory declines, our results indicate that somatosensory attenuation and precision might not be strictly linked in elderly individuals, in line with (Parthasharathy et al. 2022) but in contrast to (Wolpe et al. 2016).

## Conclusions

Overall, the results of our preregistered study suggest that aging exerts a limited influence on the perception of self-generated and externally generated touch. First, using a force-discrimination task, we observed significant somatosensory attenuation in 94% of our sample, regardless of age, extending previous findings by using different psychophysical measures (Parthasharathy et al. 2022; Wolpe et al. 2016). Second, contrary to the two preceding studies, we did not find increased attenuation in our elderly group (aged 65–77 years); however, we observed a tendency for increased attenuation when we compared the oldest subset of our elderly group (aged ≥69 years) to the young group (aged 21–33 years). Hence, our findings suggest that an increase in somatosensory attenuation might be more pronounced in samples older than 70 years. Last, we did not find an age-related decline in somatosensory precision or any indication that such a decline is related to increased somatosensory attenuation. This finding calls into question whether deficits in somatosensory precision play an important role in the age-related increase in somatosensory attenuation, as previously suggested.

## Author Contributions

All authors contributed to conceiving and designing the experiment. L.T. collected the data. L.T. and K.K. conducted the statistical analysis. L.T., X.J., K.K. and J.-J.O. wrote the manuscript.

## Acknowledgments

We thank Nora Backlund for assistance during data collection. Lili Timar and Konstantina Kilteni were supported by the Swedish Research Council (VR Starting Grant 2019-01909 to K.K.). Experiment costs were covered by the same project and the Åke Wibergs Foundation (M20-0038 granted to K.K.). Xavier Job was supported by the Swedish Research Council (VR Starting Grant 2019-01909 granted to K.K.) and the Marie Sklodowska-Curie individual fellowship (number 101059348 granted to X.J).

## Supplementary Material

Supplementary Figs. S1–S7, Texts S1-S3: https://doi.org/10.6084/m9.figshare.23904318.v1

## Disclosures

The authors declare that they have no conflicts of interest, financial or otherwise.

## Notes

### Competing Interest Statement

The authors have declared no competing interest.

### Summary of Updates

Major changes: --------------- 1) We changed all one-tailed statistical comparisons to two-tailed ones. All our conclusions remained the same. We further ameliorated the text, updated Figure 5 to describe the statistical tendencies, and improved the text. 2) We included two new paragraphs in the discussion to discuss methodological differences between the force-matching and the force-discrimination tasks. 3) We conducted additional analyses regarding the influence of working memory and all our analyses corroborated that working memory did not influence the somatosensory attenuation or precision (supplementary text S3).

