## Supplementary Material for "Aging exerts a limited influence on the perception of self-generated and externally generated touch"

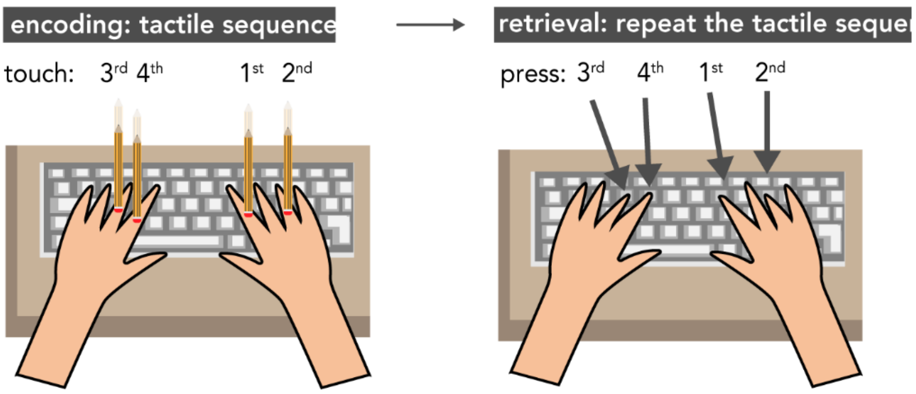


**Figure S1. The tactile working memory task.** During the task, the participant's vision was occluded. The experimenter administered a sequence of tactile stimuli (with a pencil) to the participant's fingers, (between the second and third knuckles), which were resting on eight keys of a keyboard (*left*). After the sequence was complete, the participant was asked to recall the sequence by pressing the corresponding keys in the correct order (*right*). An example trial of a 4-finger sequence is shown.

**Text S1. Cognitive function and working memory of participants**

The *elderly* group had an average MoCA score of 27.19 ± 1.17, indicating no mild cognitive decline but showed a significantly worse tactile working memory performance compared to the other two groups, consistently with previous studies showing a negative impact of age on working memory (Klencklen et al. 2017; Pliatsikas et al. 2019). Specifically, both measures of working memory were significantly lower in the *elderly* group compared to the *young* group (*longest sequence recalled*: $W = 257$, $p < .001$ *FDR-corrected*, $CI^{95} = [-2.00, -1.00]$, *rrb* = -0.603; *maximum WM score*: $t(59.6) = -5.75$, $p < .001$ *FDR-corrected*, $CI^{95} = [-5.278, -2.555]$, $d = -1.356$) and *middle-aged* group (*longest sequence recalled*: $W = 391$, $p = 0.005$ *FDR-corrected*, $CI^{95} = [-2, -0.001]$, *rrb* = -0.397; *maximum WM score*: $t(70) = -3.78$, $p < .001$ *FDR-corrected*, $CI^{95} = [-3.436, -1.064]$, $d = -0.892$) (**Figure S2a-b**). *Middle-aged* adults recalled sequences of similar length as those of *young* participants (*longest sequence recalled*: $W = 508.5$, $p = 0.108$ *FDR-corrected*, $CI^{95} = [-1, 0.001]$, *rrb* = -0.215), but they had a significantly lower maximum WM score (*maximum WM score*: $t(70) = -2.254$, $p = 0.027$ *FDR-corrected*, $CI^{95} = [-3.141, -0.192]$, $d = -0.531$).

Importantly, despite the differences among age groups, all our participants recalled sequences of at least two tactile stimuli (*longest sequence recalled* ≥ 2), which met the requirements of the force-discrimination task.

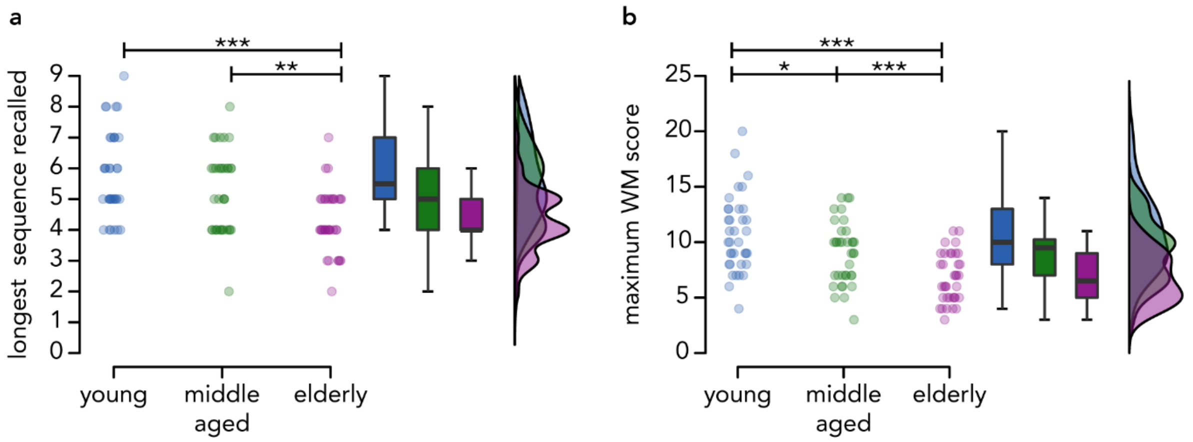


**Figure S2. Tactile working memory performance across the three age groups.** Both the *longest sequence recalled* **(a)** and the *maximum WM score* **(b)** were significantly lower in *elderly* participants than in *middle-aged* and *young* participants. Boxplots show the medians and interquartile ranges. Dot plots represent individual participant values and the raincloud plots represent the distribution of the data (*** $p$ <.001, ** $p$ <.01, * $p$ <.05). Individual data points have been horizontally jittered to avoid complete overlap.

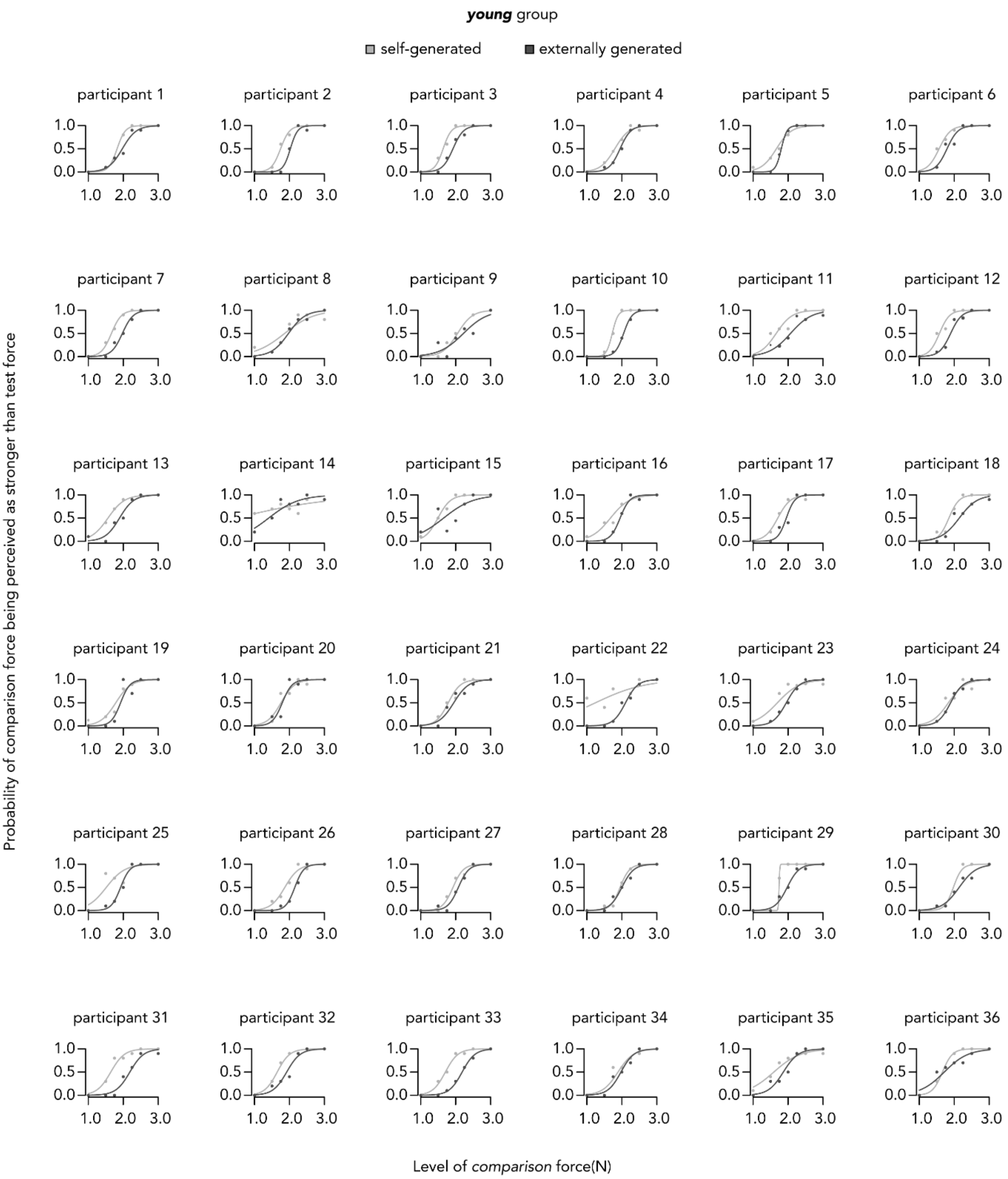


**Figure S3. Fitted logistic models of responses from *young* participants based on their responses under each condition.**

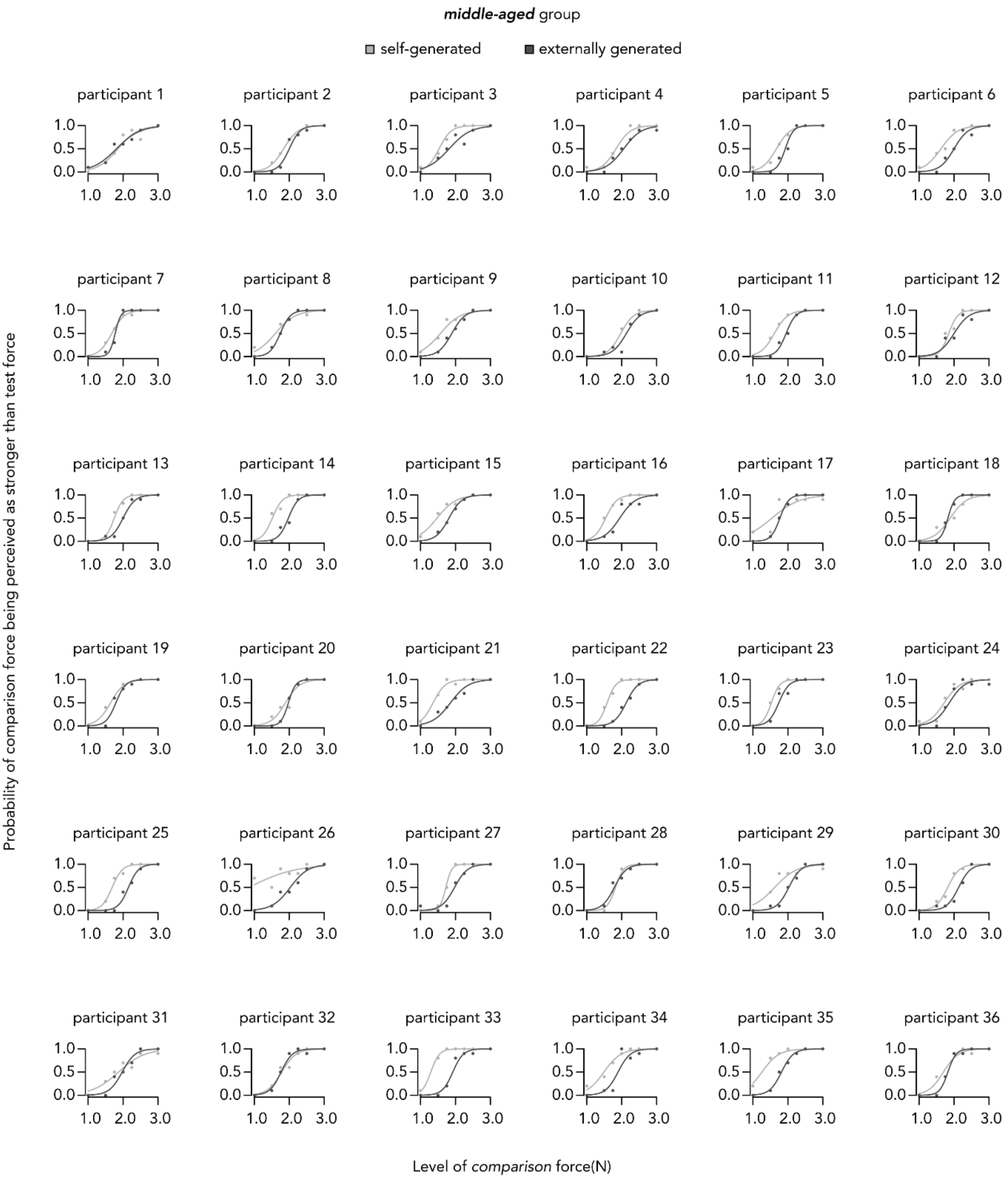


**Figure S4. Fitted logistic models of responses from *middle-aged* participants based on their responses under each condition.**

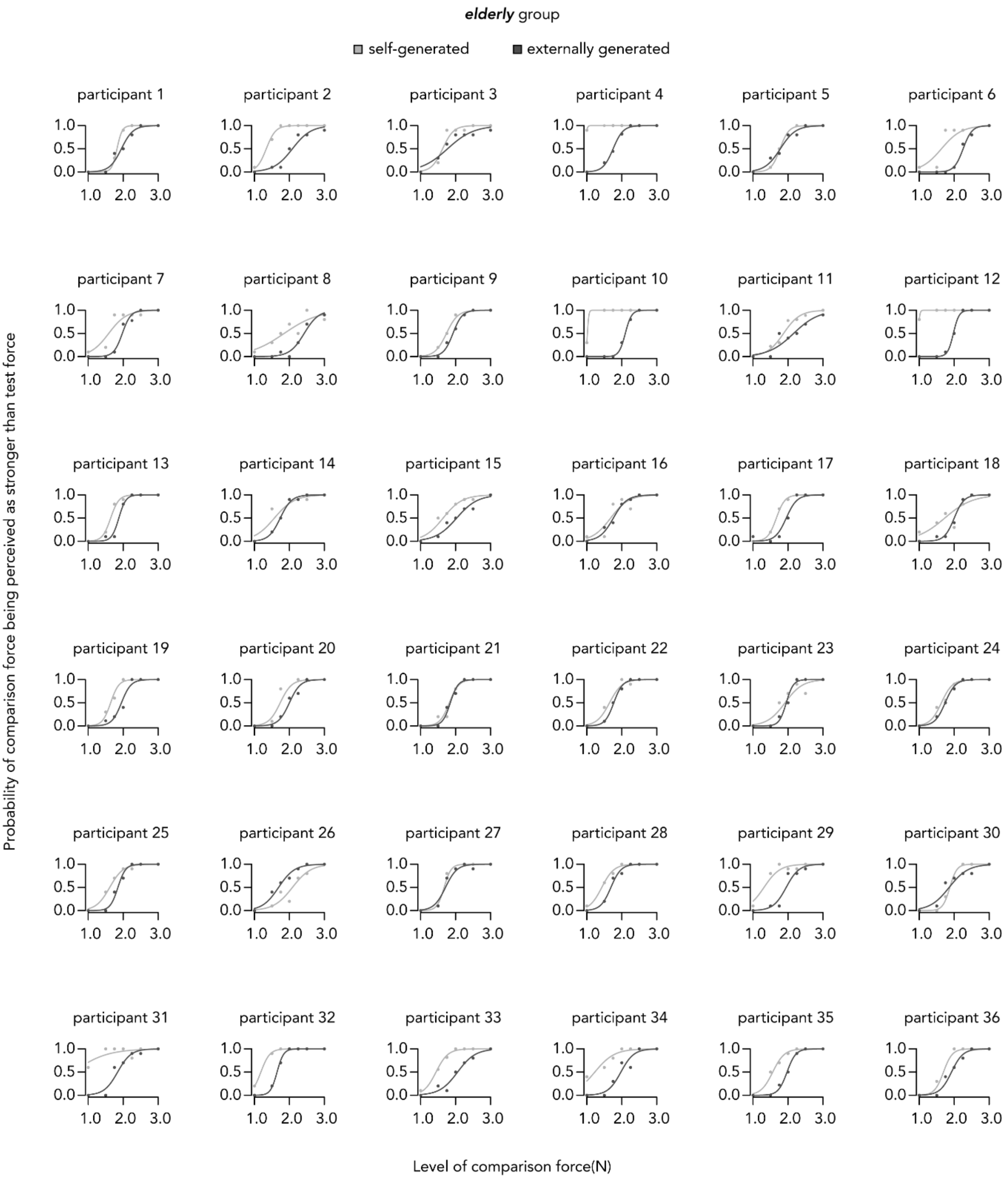


**Figure S5. Fitted logistic models of responses from *elderly* participants based on their responses under each condition.**

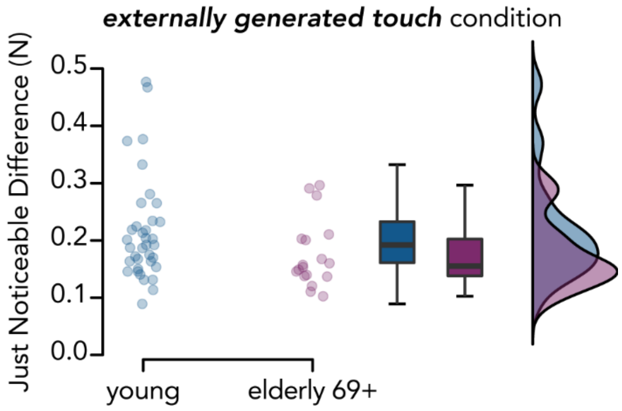


**Figure S6. Comparison of somatosensory precision between *young* (n=36) and *elderly 69+* (n=18) participants:** JNDs in the *externally generated touch* condition were not significantly different in the *elderly 69+* than in the *young* group. The boxplots display the median and interquartile ranges, and dots represent the individual participant values. Raincloud plots show the distribution of the data.

**Text S2. Active forces and times.**

Regarding the active forces, the *young* group pressed the force sensor in the *self-generated touch* condition with an average (mean ± SD) of 1.454 ± 0.648 N, the *middle-aged* pressed 1.376 ± 0.550 N and the *elderly* group pressed with an average of 1.162 ± 0.510 N. The age groups did not significantly differ in the magnitude of these forces: *elderly* vs. *young*, $W = 478$, $p = 0.156$ *FDR-corrected*, $CI^{95}$ =[-0.536, 0.006], $rrb = -0.262$, $BF_{01} = 0.676$; *elderly* vs. *middle-aged*, $W = 503$, $p = 0.156$ *FDR-corrected*, $CI^{95}$ = [-0.444, 0.042], $rrb = -0.224$, $BF_{01} = 1.011$; *middle-aged* vs. *young*, $W = 616$, $p = 0.724$ *FDR-corrected*, $CI^{95}$ = [-0.323, 0.203], $rrb$ = -0.049, $BF_{01} = 3.730$ (**Figure S7a**).

Regarding the times, the *young* group took 525.50 ± 156.38 ms on average to press on the sensor, the *middle-aged* took 504.74 ± 120.30 ms and the *elderly* group took 570.45 ± 168.22. There were no significant differences among the groups: *elderly* vs. *young*, $W = 758$, $p = 0.328$ *FDR-corrected*, $CI^{95}$ =[-28.956, 115.420], $rrb = 0.170$, $BF_{01} = 2.134$; *elderly* vs. *middle-aged*, $W = 797$, $p = 0.285$ *FDR-corrected*, $CI^{95}$ = [-8.805, 127.087], $rrb = 0.230$, $BF_{01} = 0.853$; *middle-aged* vs. *young*, $W = 631$, $p = 0.853$ *FDR-corrected*, $CI^{95}$ = [-68.716, 56.649], $rrb$ = -0.026, $BF_{01} = 3.875$ (**Figure S7b**).

We next tested for correlations between the active forces and the somatosensory attenuation and somatosensory precision. Neither the peak active force nor its time was significantly correlated with somatosensory attenuation (*peak force*, Spearman's $rho = 0.111$, $p = 0.254$, $BF_{01} = 3.639$; *time to peak force*, Spearman's $rho = -0.044$, $p = 0.651$, $BF_{01} = 8.222$) or somatosensory precision (*externally generated touch* condition: *peak force*, Spearman's $rho$ = -0.026, $p = 0.787$, $BF_{01} = 7.837$; *time to peak force*, Spearman's $rho = 0.065$, $p = 0.506$, $BF_{01}$ = 6.332; *self-generated touch* condition: *peak force*, Spearman's $rho = 0.149$, $p = 0.124$, $BF_{01}$ = 5. 296; *time to peak force*, Spearman's $rho = -0.026$, $p = 0.791$, $BF_{01} = 8.3$). To further explore whether the *peak force* could influence the magnitude of somatosensory attenuation, we performed a robust regression analysis using the *peak force* as well as the age group (*young*, *middle-aged*, *elderly*) and their interaction as regressors on somatosensory attenuation. We found no significant main effects nor interactions, suggesting that the *peak force* is not a predictor of somatosensory attenuation (all $p$-values > 0.299, $R^2 = 0.025$).

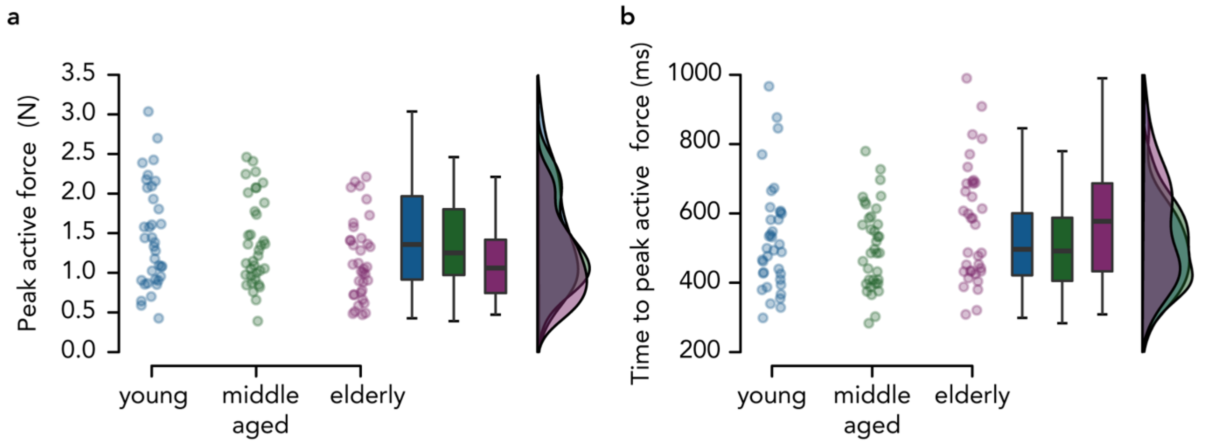

**Figure S7. Active forces and times across the three age groups.** There were no significant differences in peak active forces or the time to reach the peak active forces between the three age groups. The boxplots display the median and interquartile ranges, and dots represent the individual participant values. Raincloud plots show the distribution of the data.

**Text S3. Working memory and somatosensory attenuation.**

As mentioned above, all our participants could keep in their working memory (WM) two tactile stimuli applied on their fingers (**Supplementary Text S1**, **Figure S2**). To explore whether the working memory performance had any influence on somatosensory attenuation, we correlated both WM scores (longest sequence recalled and maximum WM score) with the PSEs in each condition separately ($PSE_{self}$ and $PSE_{external}$) as well as with somatosensory attenuation (SA). We found no significant relationship between WM and PSEs/somatosensory attenuation, and the Bayesian analyses clearly supported the absence of such relationship; somatosensory attenuation (*longest sequence recalled*: Spearman's $rho$ = 0.13, $p$ = 0.181, $BF_{01}$ = 3.088; *maximum WM score*: Spearman's $rho$ = 0.085, $p$ = 0.383, $BF_{01}$ = 5.843), $PSE_{self}$ (*longest sequence recalled*: Spearman's $rho$ = -0.138, $p$ = 0.154, $BF_{01}$ = 3.138; *maximum WM score*: Spearman's $rho$ = -0.122, $p$ = 0.210, $BF_{01}$ = 4.779), and $PSE_{external}$ (*longest sequence recalled*: Spearman's $rho$ = 0.009, $p$ = 0.929, $BF_{01}$ = 8.292; *maximum WM score*: Spearman's $rho$ = -0.076, $p$ = 0.432, $BF_{01}$ = 5.903).

In addition, we tested the WM scores as single predictors of the magnitude of somatosensory attenuation, or in addition to the age group or somatosensory precision predictors. None of the regression models yielded significant coefficients for the WM scores (all $p$-values > 0.272), suggesting that working memory had little, if any, influence on somatosensory attenuation.
